# Towards foundation models that learn across biological scales

**DOI:** 10.1101/2025.05.16.653447

**Authors:** Jeremie Kalfon, Laura Cantini, Gabriel Peyre

## Abstract

We have reached a point where many bio foundation models exist across 4 different scales, from molecules to molecular chains, cells, and tissues. However, while related in many ways, these models do not yet bridge these scales. We present a framework and architecture called Xpressor that enables cross-scale learning by (1) using a novel cross-attention mechanism to compress high-dimensional gene representations into lower-dimensional cell-state vectors, and (2) implementing a multi-scale fine-tuning approach that allows cell models to leverage and adapt protein-level representations. Using a cell Foundation Model as an example, we demonstrate that our architecture improves model performance across multiple tasks, including cell-type prediction (+12%) and embedding quality (+8%). Together, these advances represent first steps toward models that can understand and bridge different scales of biological organization.

## 1. Introduction

Biology processes information across different scales, from individual molecules to entire tissues. Recent advances in artificial intelligence have led to the development of foundation models that excel at representing biological data at specific scales, such as protein structures (Rao et al., 2020) or cell states (Kalfon et al., 2024; Cui et al., 2024). However, these models typically operate in isolation, unable to leverage the rich interconnections between different biological scales. Having models that can learn across biological scales will be crucial to capture the complexity of the biological phenomena.

The main premise of our work is that by using information gained from a lower scale (e.g., molecules), we might improve the input representations of an higher scale phenomena (e.g., cells) (Bunne et al., 2024; Song et al.). Reciprocally, using relationships learned at the higher scale, we might improve the lower-scale models too. Finally, we would want to use joint representations of molecules, DNA, proteins, cells, and tissues, which are all the elements of the organisms we want to study.

While it is likely infeasible to learn across all scales at once, we might be able to use foundation models that have been trained at specific scales, which we call uniscale models, using only fine-tuning and some architectural changes (see Figure 1). We first review the existing uniscale foundation models in depth for each of the four main biological modalities (Si et al., 2024).

**Figure 1.**
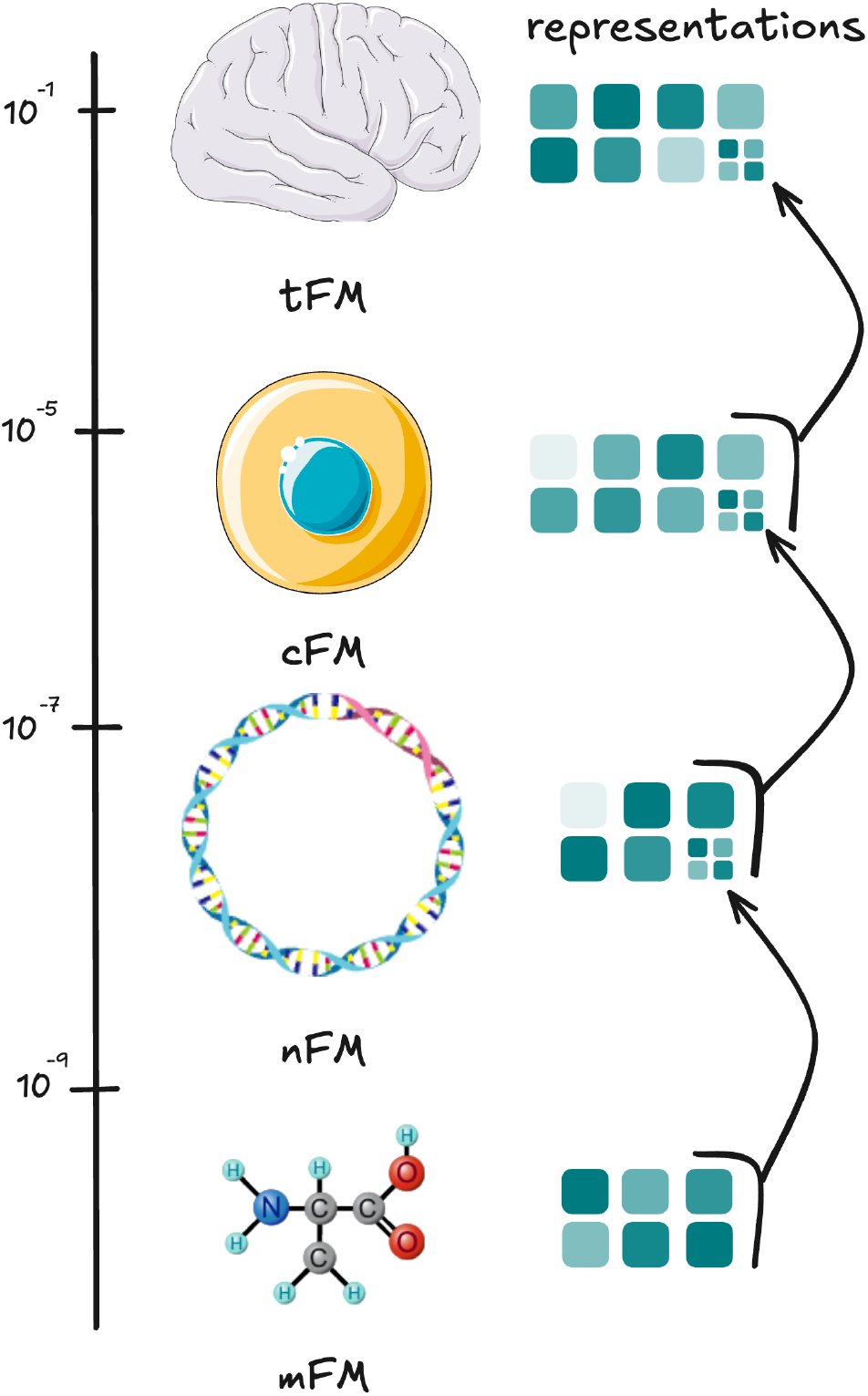
Representation of the different biological scales and how the representation of different foundation models could feed the upper scales and their learning could inform the lower scales’ representations.

### 1.1. Foundation models across scales

**Molecular foundation models (mFM)** try to model with atomistic precision the complex quantum physics-based rules that govern molecules and their interactions (Abramson et al., 2024). They generate embeddings of molecules by encoding their chemical representation, often using SMILES notation. These embeddings should contain information to predict molecular measurements such as binding to a target, potency, solubility, and more (Méndez-Lucio et al., 2024; Ross et al., 2022). The models are often built with invariances concerning the symmetries of molecules (relative positions and angles) (Batzner et al., 2022). These models can also be paired with ones that learn to predict the structure and dynamics of these molecules. Training data in this context is mostly limited by compute since molecular dynamics simulations can be generated at will (Kozinsky et al., 2023). The first use cases of such models are in material generation and drug discovery.

However, computing binding affinities and force-fields similar to the most precise molecular dynamics methods remains a frontier (Benali et al., 2025; Rhodes et al., 2025).

**Nucleotide foundation models (nFM)** are a category of models designed to analyze sequences of nucleotides or amino acids, which are encoded in triplets of nucleotides, primarily using data derived from sequencing across various life forms. Although new architectures have been introduced to handle large context sizes (Nguyen et al., 2023), most models generally rely on traditional transformer models with small context sizes and are trained with masking. These models are based on the transformer architecture and language model techniques (LM) (Vaswani et al., 2023) to produce representations of the lengthy and repetitive molecular structures found in DNA and RNA, sometimes termed dnaLM and rnaLM (Dalla-Torre et al., 2024; Wang et al., 2024a; Fradkin et al., 2024; Brixi et al., 2025).

While protein language models like ESM2 (Rao et al., 2020) have shown real-world usage in helping generate 3D models of proteins, dnaLM mainly focused on the task of understanding regulatory mechanisms, such as binding interactions and chemical modifications on DNA. It has been shown however, that representations learned by dnaLM can also contain information about the secondary structures of proteins and even protein-protein interactions (Brixi et al., 2025; Cornman et al., 2024).

For these reasons, we fold protein language models into the nFM category, proposing that their distinctions will blur in the future.

Numerous challenges still exist in accurately predicting the diverse conformations of RNA, DNA, and proteins, as well as in modeling their intricate interactions (Abramson et al., 2024). Indeed, it is still hard to measure complexes with the same accuracy as individual proteins. A goal would be to generate nFMs that learn across the very related lexicons, which are DNA, RNA, and proteins, by introducing architectures and training modalities that go beyond what exists today (Xia et al., 2025). Indeed, there we could use the framework of “learning across scales” by using the representations of molecules, learned and compressed by mFMs, as the very tokens of nFMs, allowing them to talk about ribonucleotides, deoxyribonucleotides, amino acids, and their potential modifications.

Currently, the main applications of nFMs have been in drug, and target discovery, as well as many other fields of biology.

**Cell foundation models (cFM)** are a class of models trained on a matrix of abundances of the different chemical elements (proteins, RNAs) present in cells. (Bunne et al., 2024; Kalfon et al., 2024; Theodoris et al., 2023; Cui et al., 2024; Hao et al., 2024; Rosen et al., 2023). Their architecture is often based on bidirectional encoder-based transformers trained on single-cell RNA-sequencing data. While diverse training strategies have been presented, the model’s architectures have, for now, remained fairly classical. The goal of these cFMs is to generate an accurate model of the cell that would allow predictions of cell evolution and response to perturbations (Kedzierska et al., 2023).

However, immense challenges remain. Current promises have not stood up to experimental validations (Bendidi et al.,2024; Boiarsky et al., 2023). While many reasons can be formulated, issues exist around data quality, diversity, and coverage. Indeed, single-cell data is very noisy, only measures a tiny fraction of the molecular composition of cells, and has been mostly produced on human and model animals (Program et al., 2023). While data will remain an important challenge, an area of improvement would be to, again, distill the rules of molecular interactions from sequence learned at the sequence level onto cFMs. This allows them to better learn the complex regulatory mechanisms of the cell.

**Tissue foundation models (tFM)** strive to understand the interactions between cells that form tissues, mostly in higher-order organisms. Often based on imaging techniques, they consider the 2D structural relationship of cells or group of cells in a tissue slice. The stained microscopy slides allow the prediction of tissue type, organs, and even some protein expression levels. These models are often versions of the famous vision transformer architecture and framework (Dino V2), applied to medical images (Oquab et al., 2024; Wang et al., 2024c). They thus learn on image patches where each pixel has some channels of information (often from 2 to 30 different chemical elements are represented within these channels) (Bray et al., 2016; Wenckstern et al., 2025). The number of channels can go up to tens of thousands in spatial transcriptomics image modalities, where each channel represent a transcripts location at a subcellular level(e.g., xenium) or at a cell-group-level (e.g., visium).

Overall, even more challenges arise in tissue foundation models. Most of the data exists behind institutional barriers, the resolution of high channel count modalities is really poor, while the channel amount of high-resolution modalities is really small, making it hard to predict even the cell state. Slices are often of tiny subparts of tissues. Most of the available data is in 2D slides, and 3D modalities are still burgeoning (Alon et al., 2021). We lack good measurements of what cells are communicating, but we know that they do, from sequences to molecules and even entire organelles (Hertle et al., 2021). tFMs’ vocabulary can be seen as made of cells. Their tokens are cell representations and could be be the rich representations learned by cFMs. The goal of a tFM is then to predict the presence of cells given other cells in spatial context.

### 1.2. Architectural modifications: compressed representations

For biological representations, previous methods have leveraged many different methods from matrix factorization, nearest neighbors, and neural networks (Gunawan et al., 2023; Bengio et al., 2014) amongst which popular approaches are Variational Autoencoder (VAE) such as scVI and scArches (Lopez et al., 2018; Lotfollahi et al., 2022).

In the domain of protein embedding, the HourGlass embedding method (Lu et al., 2024) introduced FSQ (Mentzer et al., 2023) as a framework to encode both amino acid sequences and 3D structural information from a pLM into a quantized latent space. Meanwhile, DNA sequence model embeddings have been mostly restricted to metagenomics, with the exception of DNA-BERT-S (Zhou et al., 2024).

Finally, it has been shown not only in biology but also in the NLP community that for transformer models, embeddings based on average,max,sum-pooling of last-layer tokens are very restrictive and do not perform well (Schockaert, 2023; Lee et al., 2025; Ilse et al., 2018). Indeed, current state-of-the-art methods use more complex approaches such as cross-attention mechanism and additional pre-training or fine-tuning tasks.

In the following, we will show that we can use a similar cross-attention mechanisms to compress the output embeddings of a foundation model into a set of lower-dimensional vectors.

### 1.3. Training modifications: fine-tuning

An extensive literature exists on fine-tuning. The simplest and most powerful approach remains to continue training on a small set of epochs and with a lower learning rate (Christophe et al., 2024). Common tools include low-rank approximations of the Multi-layer perception (MLP) and Query, Key, Values (QKV) matrices using LoRA, QLORA, and COLA (Hu et al., 2021; Dettmers et al., 2023; Ray et al., 2023; Tang et al., 2024), which allow cheap fine-tuning of large foundation models. Other common approaches also mostly revolve around reducing the memory footprint of fine-tuning by only back-propagating the loss across a specific subset of parameters, from updating only specific layers of the model, only the MLPs, the QKV matrices, or only the biases of the MLPs (Peters et al., 2019; Chronopoulou et al., 2019). Finally, adapter layers have also been used for their versatility. They often consist of an additional MLP on top of the large model’s output representations (Houlsby et al., 2019).

In the following, we will show that the adapter layer is a sensible approach to perform multi-scale fine-tuning.

### 1.4. Contributions

Following up on these recent advances, we propose:

- A cross-attention “compressor” block whose goal is to compress a foundation model’s output embeddings into a small set of low-dimensional vectors, called the *Xpressor* (Cross-Attention Compressor transformer). This is learnt using an auto-encoding approach with a reconstruction loss. The Xpressor is modality agnostic and can be used by mFMs, nFMs, cFMs, tFMs, or even other non-biological domains, and can work in addition to other training tasks like masking or denoising (see Figure 2A).

**Figure 2.**
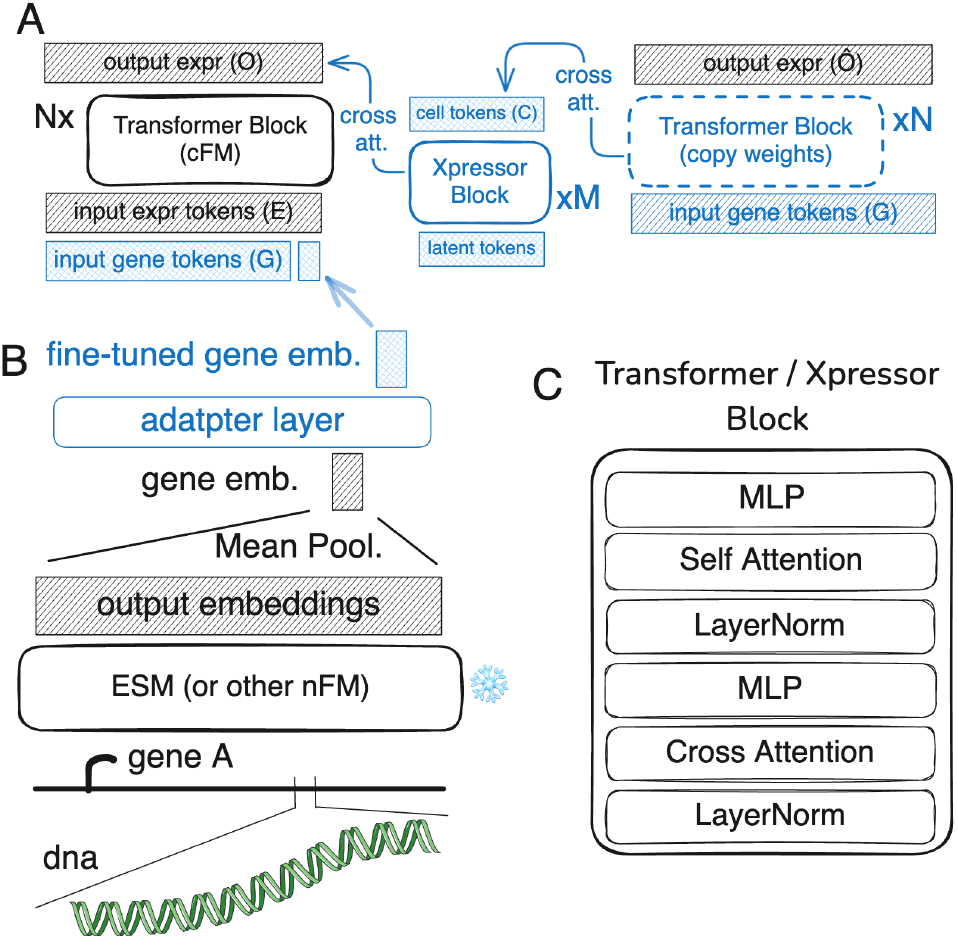
Overview of the Xpressor architecture and multi-scale fine-tuning approach applied to a cell foundation model. A. The Xpressor architecture, composed of M layers, shows how gene-level representations are compressed into cell-state vectors through cross-attention over the output embeddings of a transformer, composed of N layers. These compressed representations are then decompressed back using the same initial transformer model with cross-attention given the initial gene-level tokens. B. Example of the multi-scale fine-tuning setup illustrating how the adapter layer enables joint training of gene-level representations that are then used by a cFM. C. Detailed structure of the transformer and Xpressor blocks showing the cross-attention and selfattention sub-blocks. Blue blocks are our contributions. Shaded blocks indicate inputs and outputs.
- A multi-scale fine-tuning approach using adapter layers. This allows the fine-tuning of models from one level using the upper-scale model’s task (see Figure 2B).

## 2. Xpressor

### 2.1. Background

scPRINT (Kalfon et al., 2024) is a foundation model trained on more than 50 million unique single-cell RNA-seq profiles, representing around 100B tokens. It learns with a multi-task pre-training loss, allowing state-of-the-art zero-shot abilities in denoising and label prediction. scPRINT builds on previous foundation models, like scGPT (Cui et al., 2024) and scFoundation (Hao et al., 2024). It improves upon them on multiple benchmarks and is also easier to use and faster to train than many other similar models. Additionally, it comes with a gymnasium of benchmarks presented in Kalfon et al. (2024). For these reasons, we chose to use it as our cFM and the starting point for our work.

ESM2 (Rao et al., 2020) is a protein language model that learns embeddings of amino acid sequences. It has been shown to be able to learn the evolutionary constraints of proteins and to be able to predict contact maps. Models like ESMfold (Lin et al., 2022) have been created to predict a protein’s 3D structure directly from its output embeddings. It is also simple to use. For these reasons, we chose to use it as our nFM.

### 2.2. Approach

Our first contribution is the compression of output embeddings of foundation models using a transformer block and a bottleneck-learning training modality (see Supplementary Material 3.5): we call it the Xpressor (see Figure 2A). Compression / decompression is a key mechanism to transfer representations across scales (see Supplementary Material 1.1), we thus models that can compress and decompress their input into a lower-dimensional space. To do so, we introduce an additional set of transformer blocks called “Xpressor blocks”. In the context of scPRINT, these blocks represent cell features.

As inputs scPRINT continues to use a set of summed up gene expression and gene ID tokens. The first ones are generated using an MLP on each expression values of genes in a cell *j*, the other ones are generated from ESM2’s output embeddings of each gene sequenced aggregated with mean-pooling. The newly proposed Xpressor block uses as input a set of learned latent tokens ***T***. It then performs cross-attention between the last layer of the gene embeddings and the latent tokens (see Figure 2A). The goal is for the Xpressor blocks to be of smaller dimensions and context size than the main blocks, such that we end up with ***C*** _*j*_ a set of *n* tokens of dimension *d*_*t*_ generated from the encoded gene expression and ID matrices ***E*** _*j*_ and ***G***. Where ***G*** and ***E*** _*j*_ are sets of *m* tokens of size *d*_*c*_ representing the IDs of the genes and their corresponding expression in cell *j*, respectively, where *d*_*c*_ < *d*_*t*_ and *n* << *m*:

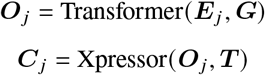

for a cell *j*, with the *Xpressor* being initialized with a learned set of input cell tokens, and ***C*** _*j*_ being the cell to-kens associated with the input ***E*** _*j*_.

The *Transformer* and *Xpressor* are both transformer with N and M layers, respectively. Indeed, we have designed both blocks to contain a cross-attention architecture (see Figure 2C) such that we can also do: ***Ô*** _*j*_ = Transformer (***C***_*j*_, ***G***), with ***Ô***_*j*_ being the output of the *Transformer* when using the *Xpressor* representation as input. We add an optional MLP after cross-attention to a transformation of the embeddings prior to the self-attention round. In our example, the decompression is done with gene ID tokens as input only (***G***) (see Figure 2A). These tokens remain the same for all cells of a given species and thus do not depend on *j*. In the context of protein language models, for example, this would be replaced by positional tokens.

As can be seen in Figure 2A, the *Transformer* blocks are applied twice. The first application act as an encoder, only using self attention, while the *Xpressor* and second application of the *Transformer* blocks act as decoders. We follow these definitions from the original “Attention is All You Need” paper (Vaswani et al., 2023). It has to be noted that in our case cross-attention is performed first instead of last. Related ideas have also been explored in Lee et al. (2025), where the authors propose a cross-attention-based method to update tokens using “latent” embeddings followed by a classical mean-pooling.

The goal of the *Xpressor* and the entire model can be seen as to perform compression of the gene tokens into a set of cell tokens similar to the classical information bottleneck from Tishby et al. (see Supplementary Material 3.5). This is our main training objective to train the *Xpressor* blocks, while the *Transformer* is also trained with masking.

In our case, each embedding represents different cell components. At training time, we present multiple losses to both regularize it and ensure differences across them, similar to what can be done in VAEs (see Supplementary Material 3.6).

### 2.3. Results

We show that such an instantiation of the transformer leads to better performance over the gymnasium of tasks available in the scPRINT cFM.

Indeed, we now look at three specific tasks: cell-type prediction, embedding quality, and gene-network inference. The tasks are the same as presented in Kalfon et al. (2024).

“Embedding quality” refers to the average scIB (Luecken et al., 2022) score for batch correction and biological consistency of cell embeddings. In this context scIB looks at the quality of the embeddings based on measures of similarity, nearest neighbors, and clustering.

Cell-label predictions are generated using a classifier on top of the cell embeddings generated by each model. We follow the approach of Kalfon et al. (2024) here, which was recently presented with a different mechanism in Wang et al. (2024b). This classification task allows us to see how one can steer the model’s embeddings to represent meaningful biological features.

Finally, we display two different metrics for gene-network inference. The gene network inference benchmark tries to estimate the quality of the self-attention matrices based on similarity to a gene-gene ground-truth matrix. Here we use EPR, an odds-ratio measure where, e.g. a value of N means that the predictions are N-times as likely to be correct as a random guess. One is the EPR score on the genome-wide perturb-seq gene-network from BenGRN (Kalfon et al., 2024), while the second is the average EPR of multiple predicted gene-networks across various cell types compared to the BenGRN’s omnipath ground truth gene network (Türei et al., 2016).

In our comparison, the regular transformer’s class-pooling is done similarly to scGPT’s (Cui et al., 2024) approach, where a class token is added to the model’s input and an additional loss is placed on it: 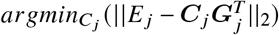. Both models use the same latent dimensions, architectures, training paradigm, and number of input tokens for both genes and cells.

We see that the Xpressor outperforms the simpler class-pooling approach on embedding quality and cell-label prediction, while the gene-network inference results remain roughly similar.

We will now see how we can further train -or fine-tune-these representations using information from the upper scale. While Xpressor layers with their small set of low-dimensional tokens are best suited for this task, we will focus on commonly available foundation models and architectures, presenting a general approach.

## 3. Multi-scale Fine-tuning

### 3.1. Background

To merge foundation models, we need a way to connect the lower-scale models to the upper one. It had been proposed in Rosen et al. (2023); Kalfon et al. (2024) to use protein language model-based representations, like those of ESM2, as input tokens for the models. This decreases the number of parameters the model has to learn; It allows the model to work on genes unseen at training time; Moreover, it also lets the model use information that it would not have gained otherwise, such as protein structure, homology, and mutations.

### 3.2. Approach

We propose going beyond simply reusing lower-scale models’ representations and fine-tuning them during the pretraining of the upper-scale model using an adapter layer (see Figure 2B). With such layer, each output embedding ***e*** is transformed with a differentiable function *f* (here, an MLP):

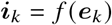

By using an MLP, the adapter layer not only applies a transformation of its input but also adds information (see Supplementary Material 3.4). In our case, we use ESM2 as the lower-scale model and scPRINT as the upper-scale model. The initial ESM2 embedding is known to contain a representation of the protein’s sequence, evolutionary similarity, and constraints.

Indeed, this is what allows this representation to replace the multiple sequence alignment (MSA) step in ESMfold (Lin et al., 2022). We posit that this initial embedding already contains the information necessary to understand some of the rules in gene interactions (homology and similar evolutionary constraints). However, representations from ESM2 are very different from those from single-cell foundation models. Our goal is to enrich these representations with knowledge gained from co-expression information across millions of cells.

### 3.3. Results

We show that a cFM trained using the pooled embeddings of a pretrained nFM performs better in most tasks from the Kalfon et al. (2024) gymnasium benchmark than one with learned representations (see Table 2). This is possible because we allow the model to start from a very rich representation instead of a random set of vectors, while still giving it the flexibility to incorporate additional knowledge. Each foundation model tested uses the same latent dimensions, architectures, training, and number of input tokens. We report the performance at the best epoch, and the training is stopped after 20 epochs.

**Table 1.** Comparison of cell embedding approaches

| Model | Cell Label<br>Pred. | Embed.<br>Quality | Gene-Net<br>Infer. |
| --- | --- | --- | --- |
| Class-pooling | 0.64 | 0.48 | **4.0,2.3** |
| **Xpressor** | **0.72** | **0.52** | 4.1,2.1 |

**Table 2.** Comparison of input-gene embedding approaches

| Model | Cell Label | Embed. | Gene-Net |
| --- | --- | --- | --- |
|  | Pred. | Quality | Infer. |
| Random init. | 0.62 | 0.48 | 4.5,1.0 |
| ESM2 frozen | 0.60 | 0.484 | 5.2,1.4 |
| **ESM2 fine-tuned** | **0.70** | **0.49** | **4.8,2.4** |

We also show the difference in cell embeddings obtained between the regular transformer and the Xpressor (see Figure 3). The dataset is a very challenging mix of modalities with various batch effects and amounts of noise. Cell types are also quite similar, making the task more difficult. We can see that the Xpressor embeddings contain more structure and resolve different cell types better than a transformer with class-pooling.

**Figure 3.**
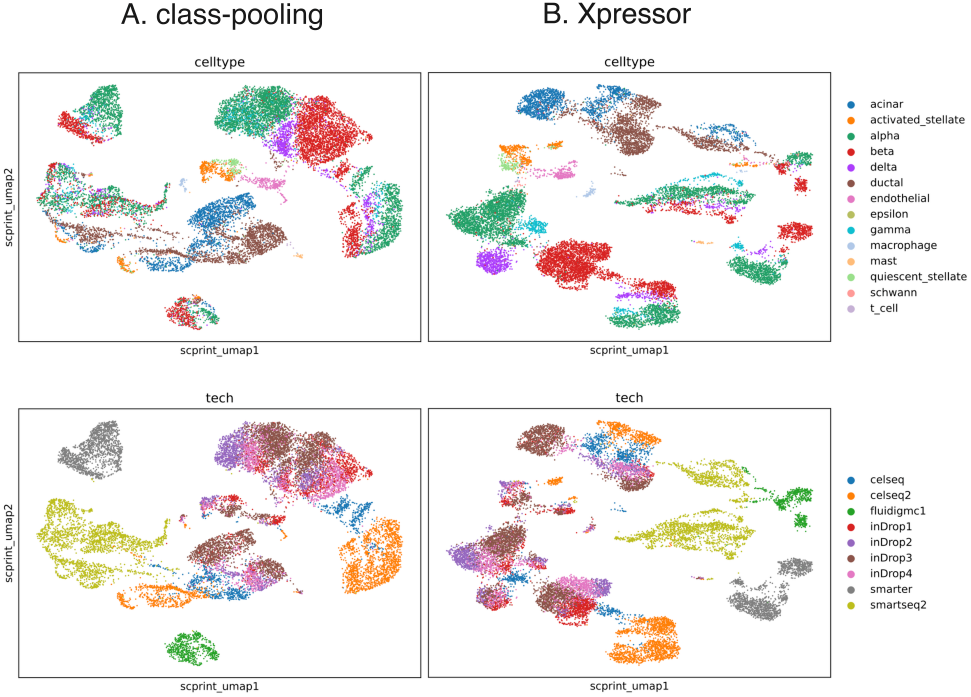
Comparison of cell embeddings between the regular transformer with class-pooling (left), scIB: 0.43, and the Xpressor (right), scIB: 0.48. The Xpressor embeddings contain more structure and resolve different cell types better.

Using ESM2’s embeddings allows scPRINT to work on genes and sequences unseen at training time, to learn from an unlimited number of species, and to integrate DNA, RNA, and protein-level information such as mutations andstructural variants.

Finally, contrary to other methods, this version does not require an update to the original model and can be added to the new model. Moreover, with this approach, scPRINT still maintains its ability to work on genes and sequences unseen at training time.

## Conclusion

We have proposed a framework towards building compositional hierarchical foundation models for life, from atoms to tissues. We highlighted progress and challenges remaining for each specific scale of biological representations. While data generation efforts focusing on breadth and quality remain paramount to progress, we believe that the composition of foundation models could drive progress forward. Having a vocabulary for biological entities will allow us to better reference them, helping us define the impact of a molecule on a tissue or the interaction between RNA and proteins. Such a model of life should not be seen as one being trained end-to-end but as a set of models distilling the key information that they have learned and that the next one requires.

We have presented one small piece in this approach, where a cell foundation model (scPRINT) uses and fine-tunes a protein sequence foundation model (ESM2). We have also shown how XPressor can compress the output representations of transformers into a small set of lower-dimensional vectors, bridging proteins to cells. Such an approach could be used to bridge molecules to proteins and cells to tissues by using compressed representations that are then fine-tuned. This is a promising back-bone architecture for a general model going from atoms to tissues.

Future work should focus on using Xpressor’s representations to power upper scale models or the ability to learn a Xpressor on top of a pre-trained foundation model. The Xpressor approach could also be extended to decoder-based language models. Finally, fine-tuning using and adaptor layer suffers from a main drawback, the non-additivity of MLPs and therefore the limited use of such fine-tuned models in other contexts than for their compressed representations. Implementing intelligent GPU scheduling and using LoRA-type methods to fine-tune only XPressor blocks will allow for more complex fine-tuning in GPU-rich settings. We will need to show that this can be applied to the other scales of biological representations and generate benchmarks that better capture the diversity of real-world biological tasks across these scales.

## Supplementary

### 3.4. proof that fine-tuning ESM2 with an adapter layer is at least sufficient to learn to add co-expression information

We show below that an MLP (with at least one hidden layer and a sufficiently large number of neurons) can learn to map each of *D* input protein embeddings to an arbitrary desired output, even if that output corresponds to a unique lookup for each protein.

1. **Finite Data Interpolation:** Let the set of *D* protein embeddings be E = {**e**_1_, **e**_2_, …, **e**_*D*_} ⊂ ℝ^*D*^, and suppose that for each **e**_*k*_ we want the MLP to output **w**_*k*_ ∈ ℝ^*D*^. Because the set E is finite, it is possible to design a function that exactly maps **e**_*k*_ ↦→ **w**_*k*_ for all *k* = 1, …, *D*.
2. **Constructive Argument Using ReLU Networks:** For a ReLU-based MLP, one can construct “bump” functions that are activated only in a small neighborhood around each **e**_*k*_. For instance, one may define functions of the form **r**_*k*_ (**x**) = σ(−∥**x** − **e**_*k*_∥ + δ), where δ > 0 is chosen so that **r**_*k*_ (**e**_*k*_) > 0 and **r**_*k*_ (**x**) is nearly zero for **x** that are not close to **e**_*k*_. By associating one or more hidden neurons to each protein embedding **e**_*k*_, one can form a linear combination MLP 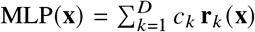, where the coefficients *c*_*k*_ ∈ ℝ^*D*^ are chosen so that MLP (**e**_*k*_) = **w**_*k*_ for all *k*. Because the supports of the functions **r**_*k*_ (**x**) can be made nearly disjoint, the MLP can “memorize” the mapping by acting as a lookup table.
3. **Conclusion:** Thus there exists a configuration of weights (and biases) in an MLP that yields MLP (**e**_*k*_) = **w**_*k*_, for *k* = 1, …, *D*. Hence, even though the MLP is simply performing a transformation, its capacity is sufficient to learn any arbitrary mapping for the *D* proteins. In other words, at worst, it can learn a mapping that is equivalent to a lookup table, thereby ensuring that each of the *D* proteins is assigned a specific, learned output value.

### 3.5. argument about the Tishby et al. bottleneck learning approach

The Information Bottleneck (IB) method seeks a stochastic mapping *p* (*t*|*x*) that compresses the input variable *X* into a representation *T*, while preserving as much information as possible about the relevant variable *Y*. The trade-off is controlled by the Lagrange multiplier *β* ≥ 0. The IB objective is to minimize the following Lagrangian:

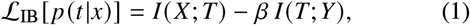

where *I* (·; ·) denotes mutual information.

Under the Markov constraint **Y** ↔ **X** ↔ **T**, the optimization leads to the following self-consistent equations:

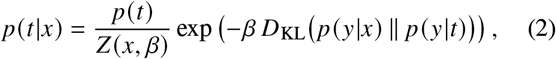

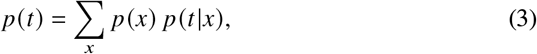

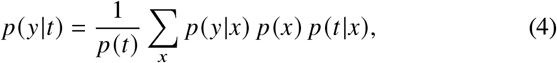

where:

- *D*_KL_ *p* (*y*|*x*) ∥ *p* (*y*|*t*) is the Kullback-Leibler divergence between the conditional distributions *p* (*y*|*x*) and *p* (*y*|*t*),
- *Z* (*x, β*) is the normalization factor ensuring that ∑_*t*_ *p* (*t* |*x*) = 1.

### 3.6. FSQ and other contrastive losses on the cell embeddings

While *D*_KL_ over a non-informative Gaussian prior is a common formulation for regularizing the embedding space in VAEs, other formulations have been used such as with the VQ-VAE and FSQ-VAE. In these contexts, the *D*_KL_ is replaced with a discretization objective tailored to the respective quantization schemes.

#### VQ-VAE

Value Quantized (VQ)-VAE employ a *codebook* of size *C*, where each codebook entry is a *d*-dimensional vector. The encoder produces a continuous latent vector, which is then mapped to its nearest entry in the codebook (a hard quantization). A commitment loss term encourages the encoder’s outputs to stay close to the chosen codebook vector, making the entire latent representation discrete at the vector level.

#### FSQ-VAE

By contrast, Finite Scalar Quantization (FSQ)-VAE discretizes each latent dimension *independently*. Specifically, the encoder outputs *d* values, each constrained to lie within a bounded range (e.g., [−1, 1]). Each dimension is then quantized into one of *M* discrete levels within that range. This dimension-wise quantization can be implemented as either a hard nearest-bin assignment or a differentiable approximation thereof. Because FSQ enforces scalar-level discretization, it provides a simpler and more fine-grained alternative to VQ’s vector-level codebook approach, while still offering strong regularization of the latent space.

#### Contrastive regularization across embedding dimensions

We further encourage each of the *d* embedding dimensions to encode distinct information by adding a contrastive loss between them. Specifically, we compute pairwise similarities among embedding elements and penalize redundancy, thus pushing each dimension to capture complementary features. A general contrast loss for this purpose can be written as 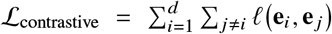, where **e**_*i*_ denotes the *i*-th embedding dimension and 𝓁 is a contrastive loss function (e.g., InfoNCE (Oord et al.)) that encourages *dissimilarity* among different embedding components.

#### Dimension-specific classifiers

To further steer each dimension’s content, one can add a separate classifier on top of each dimension to learn about different classes. The classifier for dimension *i* is trained via a cross-entropy loss 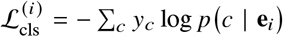, where *y*_*c*_ is the ground-truth label and *p* (*c* | **e**_*i*_) is the predicted probability for class *c*. Summing these per-dimension losses yields an overall classification objective 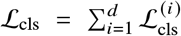. Together, the contrastive and classification losses ensure each embedding dimension captures unique, discriminative information, resulting in more expressive representations.

## Software and Data

The software and data for training scPRINT as well as gymnasium tasks and code to reproduce the results of the manuscript are available at https://github.com/cantinilab/XPressor.

WandB logs, are available in the following link: https://api.wandb.ai/links/ml4ig/h370j6io

Model checkpoints are available in the following link: https://huggingface.co/jkobject/scPRINT/tree/main

Checkpoints, wandb logs, and more will be made available after the review and deanonymization process.

## Acknowledgments

The project leading to this manuscript has received funding from the Inception program (Investissement d’Avenir grant ANR-16-CONV-0005) L.C. and the European Union (ERC StG, MULTIview-CELL, 101115618) L.C. We acknowledge the help of the HPC Core Facility of the Institut Pasteur and Déborah Philipps for the administrative support. L.C.

The work of G. Peyré was supported by the French government under management of Agence Nationale de la Recherche as part of the ‘Investissements d’avenir’ program, reference ANR19-P3IA-0001 (PRAIRIE 3IA Institute). G.P.

## Impact Statement

This paper presents work whose goal is to advance the fields of computational biology and machine learning. No ethical issues are raised by the work other than what is typically noted in computational biology and foundation model papers. It might have an impact on building better models for drug discovery, target discovery, and improving our understanding of biological systems.

## References

Abramson, J., Adler, J., Dunger, J., Evans, R., Green, T., Pritzel, A., Ronneberger, O., Willmore, L., Ballard, A. J., Bambrick, J., Bodenstein, S. W., Evans, D. A., Hung, C.-C., O’Neill, M., Reiman, D., Tunyasuvunakool, K., Wu, Z., Žemgulytė, A., Arvaniti, E., Beattie, C., Bertolli, O., Bridgland, A., Cherepanov, A., Congreve, M., Cowen-Rivers, A. I., Cowie, A., Figurnov, M., Fuchs, F. B., Gladman, H., Jain, R., Khan, Y. A., Low, C. M. R., Perlin, K., Potapenko, A., Savy, P., Singh, S., Stecula, A., Thillaisundaram, A., Tong, C., Yakneen, S., Zhong, E. D., Zielinski, M., Žídek, A., Bapst, V., Kohli, P., Jaderberg, M., Hassabis, D., and Jumper, J. M. Accurate structure prediction of biomolecular interactions with AlphaFold 3. Nature, 630(8016):493–500, June 2024. ISSN 1476-4687. doi: 10.1038/s41586-024-07487-w.

Alon, S., Goodwin, D. R., Sinha, A., Wassie, A. T., Chen, F., Daugharthy, E. R., Bando, Y., Kajita, A., Xue, A. G., Marrett, K., Prior, R., Cui, Y., Payne, A. C., Yao, C.-C., Suk, H.-J., Wang, R., Yu, C.-C. J., Tillberg, P., Reginato, P., Pak, N., Liu, S., Punthambaker, S., Iyer, E. P. R., Kohman, R. E., Miller, J. A., Lein, E. S., Lako, A., Cullen, N., Rodig, S., Helvie, K., Abravanel, D. L., Wagle, N., Johnson, B. E., Klughammer, J., Slyper, M., Waldman, J., Jané-Valbuena, J., Rozenblatt-Rosen, O., Regev, A., Imaxt Consortium, Church, G. M., Marblestone, A. H., and Boyden, E. S. Expansion sequencing: Spatially precise in situ transcriptomics in intact biological systems. Science, 371(6528):eaax2656, January 2021. doi: 10.1126/science.aax2656.

Batzner, S., Musaelian, A., Sun, L., Geiger, M., Mailoa, J. P., Kornbluth, M., Molinari, N., Smidt, T. E., and Kozinsky, B. E(3)-equivariant graph neural networks for data-efficient and accurate interatomic potentials. Nature Communications, 13(1):2453, May 2022. ISSN 2041-1723. doi: 10.1038/s41467-022-29939-5.

Benali, A., Plé, T., Adjoua, O., Agarawal, V., Applencourt, T., Blazhynska, M., Iii, R. C., Gasperich, K., Hossain, K., Kim, J., Knight, C., Krogel, J. T., Maday, Y., Maria, M., Montes, M., Luo, Y., Posenitskiy, E., Villot, C., Vishwanath, V., Lagardère, L., and Piquemal, J.-P. Pushing the accuracy limit of foundation neural network models with quantum monte carlo forces and path integrals, 2025. URL https://arxiv.org/abs/2504.07948.

Bendidi, I., Whitfield, S., Kenyon-Dean, K., Yedder, H. B., Mesbahi, Y. E., Noutahi, E., and Denton, A. K. Bench-marking Transcriptomics Foundation Models for Perturbation Analysis : One PCA still rules them all, November 2024.

Bengio, Y., Courville, A., and Vincent, P. Representation Learning: A Review and New Perspectives, April 2014.

Boiarsky, R., Singh, N., Buendia, A., Getz, G., and Sontag, D. A Deep Dive into Single-Cell RNA Sequencing Foundation Models, October 2023.

Bray, M.-A., Singh, S., Han, H., Davis, C. T., Borgeson, B., Hartland, C., Kost-Alimova, M., Gustafsdottir, S. M., Gibson, C. C., and Carpenter, A. E. Cell Painting, a high-content image-based assay for morphological profiling using multiplexed fluorescent dyes. Nature Protocols, 11 (9):1757–1774, September 2016. ISSN 1750-2799. doi: 10.1038/nprot.2016.105.

Brixi, G., Durrant, M. G., Ku, J., Poli, M., Brockman, G., Chang, D., Gonzalez, G. A., King, S. H., Li, D. B., Merchant, A. T., Naghipourfar, M., Nguyen, E., Ricci-Tam, C., Romero, D. W., Sun, G., Taghibakshi, A., Vorontsov, A., Yang, B., Deng, M., Gorton, L., Nguyen, N., Wang, N. K., Adams, E., Baccus, S. A., Dillmann, S., Ermon, S., Guo, D., Ilango, R., Janik, K., Lu, A. X., Mehta, R., Mofrad, M. R. K., Ng, M. Y., Pannu, J., Ré, C., Schmok, J. C., John, J. S., Sullivan, J., Zhu, K., Zynda, G., Balsam, D., Collison, P., Costa, A. B., Hernandez-Boussard, T., Ho, E., Liu, M.-Y., McGrath, T., Powell, K., Burke, D. P., Goodarzi, H., Hsu, P. D., and Hie, B. L. Genome modeling and design across all domains of life with Evo 2, February 2025.

Bunne, C., Roohani, Y., Rosen, Y., Gupta, A., Zhang, X., Roed, M., Alexandrov, T., AlQuraishi, M., Brennan, P., Burkhardt, D. B., Califano, A., Cool, J., Dernburg, A. F., Ewing, K., Fox, E. B., Haury, M., Herr, A. E., Horvitz, E., Hsu, P. D., Jain, V., Johnson, G. R., Kalil, T., Kelley, D. R., Kelley, S. O., Kreshuk, A., Mitchison, T., Otte, S., Shendure, J., Sofroniew, N. J., Theis, F., Theodoris, C. V., Upadhyayula, S., Valer, M., Wang, B., Xing, E., Yeung-Levy, S., Zitnik, M., Karaletsos, T., Regev, A., Lundberg, E., Leskovec, J., and Quake, S. R. How to build the virtual cell with artificial intelligence: Priorities and opportunities. Cell, 187(25):7045–7063, December 2024. ISSN 0092-8674, 1097-4172. doi: 10.1016/j.cell.2024.11.015.

Christophe, C., Kanithi, P. K., Munjal, P., Raha, T., Hayat, N., Rajan, R., Al-Mahrooqi, A., Gupta, A., Salman, M. U., Gosal, G., Kanakiya, B., Chen, C., Vassilieva, N., Amor, B. B., Pimentel, M. A., and Khan, S. Med42 – Evaluating Fine-Tuning Strategies for Medical LLMs: Full-Parameter vs. Parameter-Efficient Approaches, April 2024.

Chronopoulou, A., Baziotis, C., and Potamianos, A. An Embarrassingly Simple Approach for Transfer Learning from Pretrained Language Models, May 2019.

Cornman, A., West-Roberts, J., Camargo, A. P., Roux, S., Beracochea, M., Mirdita, M., Ovchinnikov, S., and Hwang, Y. The OMG dataset: An Open MetaGenomic corpus for mixed-modality genomic language modeling, August 2024.

Cui, H., Wang, C., Maan, H., Pang, K., Luo, F., Duan, N., and Wang, B. scGPT: Toward building a foundation model for single-cell multi-omics using generative AI. Nature Methods, pp. 1–11, February 2024. ISSN 1548-7105. doi: 10.1038/s41592-024-02201-0.

Dalla-Torre, H., Gonzalez, L., Mendoza-Revilla, J., Carranza, N. L., Grzywaczewski, A. H., Oteri, F., Dallago, C., Trop, E., de Almeida, B. P., Sirelkhatim, H., Richard, G., Skwark, M., Beguir, K., Lopez, M., and Pierrot, T. The Nucleotide Transformer: Building and Evaluating Robust Foundation Models for Human Genomics, October 2024.

Dettmers, T., Pagnoni, A., Holtzman, A., and Zettlemoyer, L. QLoRA: Efficient Finetuning of Quantized LLMs, May 2023.

Fradkin, P., Shi, R., Isaev, K., Frey, B. J., Morris, Q., Lee, L. J., and Wang, B. Orthrus: Towards Evolutionary and Functional RNA Foundation Models, October 2024.

Gunawan, I., Vafaee, F., Meijering, E., and Lock, J. G. An introduction to representation learning for single-cell data analysis. Cell Reports Methods, 3(8):100547, August 2023. ISSN 2667-2375. doi: 10.1016/j.crmeth.2023.100547.

Hao, M., Gong, J., Zeng, X., Liu, C., Guo, Y., Cheng, X., Wang, T., Ma, J., Zhang, X., and Song, L. Large-scale foundation model on single-cell transcriptomics. Nature Methods, pp. 1–11, June 2024. ISSN 1548-7105. doi: 10.1038/s41592-024-02305-7.

Hertle, A. P., Haberl, B., and Bock, R. Horizontal genome transfer by cell-to-cell travel of whole organelles. Science Advances, 7(1):eabd8215, January 2021. doi: 10.1126/sciadv.abd8215.

Houlsby, N., Giurgiu, A., Jastrzebski, S., Morrone, B., de Laroussilhe, Q., Gesmundo, A., Attariyan, M., and Gelly, S. Parameter-Efficient Transfer Learning for NLP, June 2019.

Hu, E. J., Shen, Y., Wallis, P., Allen-Zhu, Z., Li, Y., Wang, S., Wang, L., and Chen, W. LoRA: Low-Rank Adaptation of Large Language Models, October 2021.

Ilse, M., Tomczak, J., and Welling, M. Attention-based Deep Multiple Instance Learning. In Proceedings of the 35th International Conference on Machine Learning, pp. 2127–2136. PMLR, July 2018.

Kalfon, J., Samaran, J., Peyré, G., and Cantini, L. scPRINT: Pre-training on 50 million cells allows robust gene network predictions, July 2024.

Kedzierska, K. Z., Crawford, L., Amini, A. P., and Lu, A. X. Assessing the limits of zero-shot foundation models in single-cell biology, October 2023.

Kozinsky, B., Musaelian, A., Johansson, A., and Batzner, S. Scaling the Leading Accuracy of Deep Equivariant Models to Biomolecular Simulations of Realistic Size. In Proceedings of the International Conference for High Performance Computing, Networking, Storage and Analysis, SC ‘23, pp. 1–12, New York, NY, USA, November 2023. Association for Computing Machinery. ISBN 9798400701092. doi: 10.1145/3581784.3627041.

Lee, C., Roy, R., Xu, M., Raiman, J., Shoeybi, M., Catanzaro, B., and Ping, W. NV-Embed: Improved Techniques for Training LLMs as Generalist Embedding Models, January 2025.

Lin, Z., Akin, H., Rao, R., Hie, B., Zhu, Z., Lu, W., Costa, A. d. S., Fazel-Zarandi, M., Sercu, T., Candido, S., and Rives, A. Language models of protein sequences at the scale of evolution enable accurate structure prediction, July 2022.

Lopez, R., Regier, J., Cole, M. B., Jordan, M. I., and Yosef, N. Deep generative modeling for single-cell transcriptomics. Nature Methods, 15(12):1053–1058, December 2018. ISSN 1548-7105. doi: 10.1038/s41592-018-0229-2.

Lotfollahi, M., Naghipourfar, M., Luecken, M. D., Khajavi, M., Büttner, M., Wagenstetter, M., Avsec, Ž., Gayoso, A., Yosef, N., Interlandi, M., Rybakov, S., Misharin, A. V., and Theis, F. J. Mapping single-cell data to reference atlases by transfer learning. Nature Biotechnology, 40 (1):121–130, January 2022. ISSN 1546-1696. doi: 10.1038/s41587-021-01001-7.

Lu, A. X., Yan, W., Yang, K. K., Gligorijevic, V., Cho, K., Abbeel, P., Bonneau, R., and Frey, N. Tokenized and Continuous Embedding Compressions of Protein Sequence and Structure, November 2024.

Luecken, M. D., Büttner, M., Chaichoompu, K., Danese, A., Interlandi, M., Mueller, M. F., Strobl, D. C., Zappia, L., Dugas, M., Colomé-Tatché, M., and Theis, F. J. Benchmarking atlas-level data integration in single-cell genomics. Nature Methods, 19(1):41–50, January 2022. ISSN 1548-7105. doi: 10.1038/s41592-021-01336-8.

Méndez-Lucio, O., Nicolaou, C. A., and Earnshaw, B. MolE: A foundation model for molecular graphs using disentangled attention. Nature Communications, 15(1): 9431, November 2024. ISSN 2041-1723. doi: 10.1038/s41467-024-53751-y.

Mentzer, F., Minnen, D., Agustsson, E., and Tschannen, M. Finite Scalar Quantization: VQ-VAE Made Simple, October 2023.

Nguyen, E., Poli, M., Faizi, M., Thomas, A., Birch-Sykes, C., Wornow, M., Patel, A., Rabideau, C., Massaroli, S., Bengio, Y., Ermon, S., Baccus, S. A., and Ré, C. Hyenadna: Long-range genomic sequence modeling at single nucleotide resolution, 2023. URL https://arxiv.org/abs/2306.15794.

Oord, A. v. d., Li, Y., and Vinyals, O. Representation Learning with Contrastive Predictive Coding. doi: 10.48550/arXiv.1807.03748.

Oquab, M., Darcet, T., Moutakanni, T., Vo, H., Szafraniec, M., Khalidov, V., Fernandez, P., Haziza, D., Massa, F., El-Nouby, A., Assran, M., Ballas, N., Galuba, W., Howes, R., Huang, P.-Y., Li, S.-W., Misra, I., Rabbat, M., Sharma, V., Synnaeve, G., Xu, H., Jegou, H., Mairal, J., Labatut, P., Joulin, A., and Bojanowski, P. DINOv2: Learning Robust Visual Features without Supervision, February 2024.

Peters, M. E., Ruder, S., and Smith, N. A. To Tune or Not to Tune? Adapting Pretrained Representations to Diverse Tasks, June 2019.

Program, C. S.-C. B., Abdulla, S., Aevermann, B., Assis, P., Badajoz, S., Bell, S. M., Bezzi, E., Cakir, B., Chaffer, J., Chambers, S., Cherry, J. M., Chi, T., Chien, J., Dorman, L., Garcia-Nieto, P., Gloria, N., Hastie, M., Hegeman, D., Hilton, J., Huang, T., Infeld, A., Istrate, A.-M., Jelic, I., Katsuya, K., Kim, Y. J., Liang, K., Lin, M., Lombardo, M., Marshall, B., Martin, B., McDade, F., Megill, C., Patel, N., Predeus, A., Raymor, B., Robatmili, B., Rogers, D., Rutherford, E., Sadgat, D., Shin, A., Small, C., Smith, T., Sridharan, P., Tarashansky, A., Tavares, N., Thomas, H., Tolopko, A., Urisko, M., Yan, J., Yeretssian, G., Zamanian, J., Mani, A., Cool, J., and Carr, A. CZ CELL×GENE Discover: A single-cell data platform for scalable exploration, analysis and modeling of aggregated data, November 2023.

Rao, R., Meier, J., Sercu, T., Ovchinnikov, S., and Rives, A. Transformer protein language models are unsupervised structure learners, December 2020.

Ray, A., Radenovic, F., Dubey, A., Plummer, B. A., Krishna, R., and Saenko, K. COLA: A Benchmark for Compositional Text-to-image Retrieval, November 2023.

Rhodes, B., Vandenhaute, S., Šimkus, V., Gin, J., Godwin, J., Duignan, T., and Neumann, M. Orb-v3: atomistic simulation at scale, 2025. URL https://arxiv.org/abs/2504.06231.

Rosen, Y., Roohani, Y., Agarwal, A., Samotorčan, L., Consortium, T. S., Quake, S. R., and Leskovec, J. Universal Cell Embeddings: A Foundation Model for Cell Biology, November 2023.

Ross, J., Belgodere, B., Chenthamarakshan, V., Padhi, I., Mroueh, Y., and Das, P. Large-scale chemical language representations capture molecular structure and properties. Nature Machine Intelligence, 4(12):1256– 1264, December 2022. ISSN 2522-5839. doi: 10.1038/s42256-022-00580-7.

Schockaert, S. Embeddings as Epistemic States: Limitations on the Use of Pooling Operators for Accumulating Knowledge, July 2023.

Si, Y., Zou, J., Gao, Y., Chuai, G., Liu, Q., and Chen, L. Foundation models in molecular biology. Biophysics Reports, 10(3):135–151, June 2024. ISSN 2364-3439. doi: 10.52601/bpr.2024.240006.

Song, L., Segal, E., and Xing, E. Toward AI-Driven Digital Organism: Multiscale Foundation Models for Predicting, Simulating and Programming Biology at All Levels. URL http://arxiv.org/abs/2412.06993.

Tang, Z., Somia, N., Yu, Y., and Koo, P. K. Evaluating the representational power of pre-trained DNA language models for regulatory genomics. bioRxiv, pp. 2024.02.29.582810, September 2024. ISSN 2692-8205. doi: 10.1101/2024.02.29.582810.

Theodoris, C. V., Xiao, L., Chopra, A., Chaffin, M. D., Al Sayed, Z. R., Hill, M. C., Mantineo, H., Brydon, E. M., Zeng, Z., Liu, X. S., and Ellinor, P. T. Transfer learning enables predictions in network biology. Nature, 618(7965):616–624, June 2023. ISSN 1476-4687. doi: 10.1038/s41586-023-06139-9.

Tishby, N., Pereira, F. C., and Bialek, W. The Information Bottleneck Method.

Türei, D., Korcsmáros, T., and Saez-Rodriguez, J. OmniPath: Guidelines and gateway for literature-curated signaling pathway resources. Nature Methods, 13(12):966– 967, December 2016. ISSN 1548-7105. doi: 10.1038/nmeth.4077.

Vaswani, A., Shazeer, N., Parmar, N., Uszkoreit, J., Jones, L., Gomez, A. N., Kaiser, L., and Polosukhin, I. Attention Is All You Need, August 2023.

Wang, N., Bian, J., Li, Y., Li, X., Mumtaz, S., Kong, L., and Xiong, H. Multi-purpose RNA language modelling with motif-aware pretraining and type-guided fine-tuning. Nature Machine Intelligence, 6(5):548–557, May 2024a. ISSN 2522-5839. doi: 10.1038/s42256-024-00836-4.

Wang, Q., Zhu, H., Hu, Y., Chen, Y., Wang, Y., Zhang, X., Zou, J., Kellis, M., Li, Y., Liu, D., and Jiang, L. Hierarchical Interpretation of Out-of-Distribution Cells Using Bottlenecked Transformer, December 2024b.

Wang, X., Zhao, J., Marostica, E., Yuan, W., Jin, J., Zhang, J., Li, R., Tang, H., Wang, K., Li, Y., Wang, F., Peng, Y., Zhu, J., Zhang, J., Jackson, C. R., Zhang, J., Dillon, D., Lin, N. U., Sholl, L., Denize, T., Meredith, D., Ligon, K. L., Signoretti, S., Ogino, S., Golden, J. A., Nasrallah, M. P., Han, X., Yang, S., and Yu, K.-H. A pathology foundation model for cancer diagnosis and prognosis prediction. Nature, 634(8035):970–978, October 2024c. ISSN 1476-4687. doi: 10.1038/s41586-024-07894-z.

Wenckstern, J., Jain, E., Vasilev, K., Pariset, M., Wicki, A., Gut, G., and Bunne, C. AI-powered virtual tissues from spatial proteomics for clinical diagnostics and biomedical discovery, January 2025.

Xia, Y., Jin, P., Xie, S., He, L., Cao, C., Luo, R., Liu, G., Wang, Y., Liu, Z., Chen, Y.-J., Guo, Z., Bai, Y., Deng, P., Min, Y., Lu, Z., Hao, H., Yang, H., Li, J., Liu, C., Zhang, J., Zhu, J., Wu, K., Zhang, W., Gao, K., Pei, Q., Wang, Q., Liu, X., Li, Y., Zhu, H., Lu, Y., Ma, M., Wang, Z., Xie, T., Maziarz, K., Segler, M., Yang, Z., Chen, Z., Shi, Y., Zheng, S., Wu, L., Hu, C., Dai, P., Liu, T.-Y., Liu, H., and Qin, T. NatureLM: Deciphering the Language of Nature for Scientific Discovery, February 2025.

Zhou, Z., Wu, W., Ho, H., Wang, J., Shi, L., Davuluri, R. V., Wang, Z., and Liu, H. DNABERT-S: Pioneering Species Differentiation with Species-Aware DNA Embeddings, October 2024.

